# InteracTor: A new integrative feature extraction toolkit for improved characterization of protein structural properties

**DOI:** 10.1101/2024.10.07.616705

**Authors:** Jose Cleydson F. Silva, Layla Schuster, Nick Sexson, Matias Kirst, Marcio F. R. Resende, Raquel Dias

**Affiliations:** Department of Microbiology and Cell Science, Institute of Food and Agricultural Sciences, University of Florida, 1052 Museum Road, 32611-0700, Gainesville, FL, USA; Horticultural Sciences Department, University of Florida, Gainesville, FL, United States; Genetics Institute, University of Florida, Gainesville, FL, USA; School of Forest, Fisheries and Geomatic Sciences, University of Florida, Gainesville, FL, USA

**Keywords:** Feature Extraction, Interatomic Interactions, Structural Diversity, structural biology, Data Visualization, Statistical Analysis

## Abstract

Understanding the structural and functional diversity of protein families is crucial for elucidating their biological roles. Traditional analyses often focus on primary and secondary structures, which include amino acid sequences and local folding patterns like alpha helices and beta sheets. However, primary and secondary structures alone may not fully represent the complex interactions within proteins. To address this limitation, we developed a new algorithm (InteracTor) to analyze proteins by extracting features from their three-dimensional (3D) structures. The toolkit extracts interatomic interaction features such as hydrogen bonds, van der Waals interactions, and hydrophobic contacts, which are crucial for understanding protein dynamics, structure, and function. Incorporating 3D structural data and interatomic interaction features provides a more comprehensive understanding of protein structure and function, potentially enhancing downstream predictive modeling capabilities. By using the extracted features in Mutual Information scoring (MI), Principal Component Analysis (PCA), t-distributed Stochastic Neighbor Embedding (t-SNE), Uniform Manifold Approximation and Projection (UMAP), and hierarchical clustering analysis as use cases, we identified clear separations among protein structural families, highlighting distinct functional aspects. Our analysis revealed that interatomic interaction features were more informative than protein secondary structure features, providing insights into potential structural and functional properties. These findings underscore the significance of considering tertiary structure in protein analysis, offering a robust framework for future studies aiming at enhancing the capabilities of models for protein function prediction and drug discovery.

## Introduction

In recent decades, high-throughput sequencing techniques have dramatically expanded protein sequence databases, while advances in cryo-electron microscopy and deep-learning-based computational structure determination methods, including AlphaFold [1] and RoseTTAFold [2], have transformed protein structure elucidation. This surge in available sequence and structural data has catalyzed the development of machine- and deep-learning techniques for predictive modeling. This data has been leveraged to address a variety of challenges, such as identifying non-classical secreted proteins [3, 4], predicting binding affinity [5–7], and engineering proteins for novel functions [8, 9].

Central to these algorithms is the selection of feature encodings, aimed at converting protein sequences and physiochemical properties into machine-readable formats. Ideally, this process captures the attributes most relevant to the predictive targets of interest. Sequence-based feature representations are among the most widely utilized, including amino acid composition, chemical property-based features, k-mers, and alignment-based embeddings. These descriptors effectively simplify sequence information and reduce the data dimensionality while still highlighting broader functional characteristics, sequence patterns, and evolutionary relationships. However, sequence-based methods can suffer from high dimensionality and data sparsity and are limited in their ability to capture critical properties influencing protein function. The incorporation of 3D structural data into the suite of available encodings allows predictive models to have a deeper layer of biological context that can give insight into functional dynamics. This has spurred the development of more comprehensive feature extraction platforms such as iFeatureOmega [10], Pfeature [11], and ProFeatX [12]. which incorporate both sequence and structural descriptors.

While these descriptors have been foundational for many types of predictive modeling, the encoding of interaction features represents a significant yet underexploited strategy. These features, including dispersion forces, Van der Waals interactions, hydrophobic interactions, and repulsive interactions, add another dimension to the protein representation. Beyond informing protein-protein and protein-ligand interactions, interaction features within the same protein can significantly enhance the capabilities of models that predict protein function. Furthermore, such interaction-based descriptions may also enhance drug-discovery capabilities and provide insights into protein dynamics, folding, and stability.

Here we present InteracTor, a new toolkit for the extraction of three types of protein feature encodings: interaction features, physicochemical features, and compositional features. Interaction features include hydrogen bonds, hydrophobic contacts, repulsive interactions, and van der Waals interactions, each encoding aspects of molecular dynamics that play an important role in governing protein function. Specifically, hydrogen bonds and hydrophobic contacts are important for stabilizing secondary and tertiary structures [13, 14]. Van der Waals interactions influence molecular complementarity, which is crucial for substrate binding, and mediate transient interactions that can facilitate or destabilize protein structures and complexes [15]. Physiochemical property features include accessible solvent area, hydrophobicity, and surface tension, which are implicated in protein folding, stability, solubility, and protein-protein interactions. Compositional features include mono-, di-, and tripeptide composition and amino acid side chain chemical property (CPAASC) frequencies [16, 17, 18]. These features determine local spatial arrangements (secondary structure) and the overall 3D folded conformation (tertiary structure) of a protein through the formation of alpha helices, beta sheets, loops, and structural motifs.

Our integrative feature encoding approach underscores the complexity and richness of proteins, pointing toward a more holistic approach in computational biology. Structural and sequence-based features are complementary and provide a more comprehensive representation of the protein, ultimately improving the performance of tasks like protein structure and function characterization.

## Materials and Methods

### Algorithm description

The InteracTor algorithm consists of a sequence of steps for analyzing protein structures and calculating various features from proteins’ interatomic interactions, physicochemical properties of residues, and peptide composition (**Figure 1**). The following is a description of each step:

**Figure 1:**
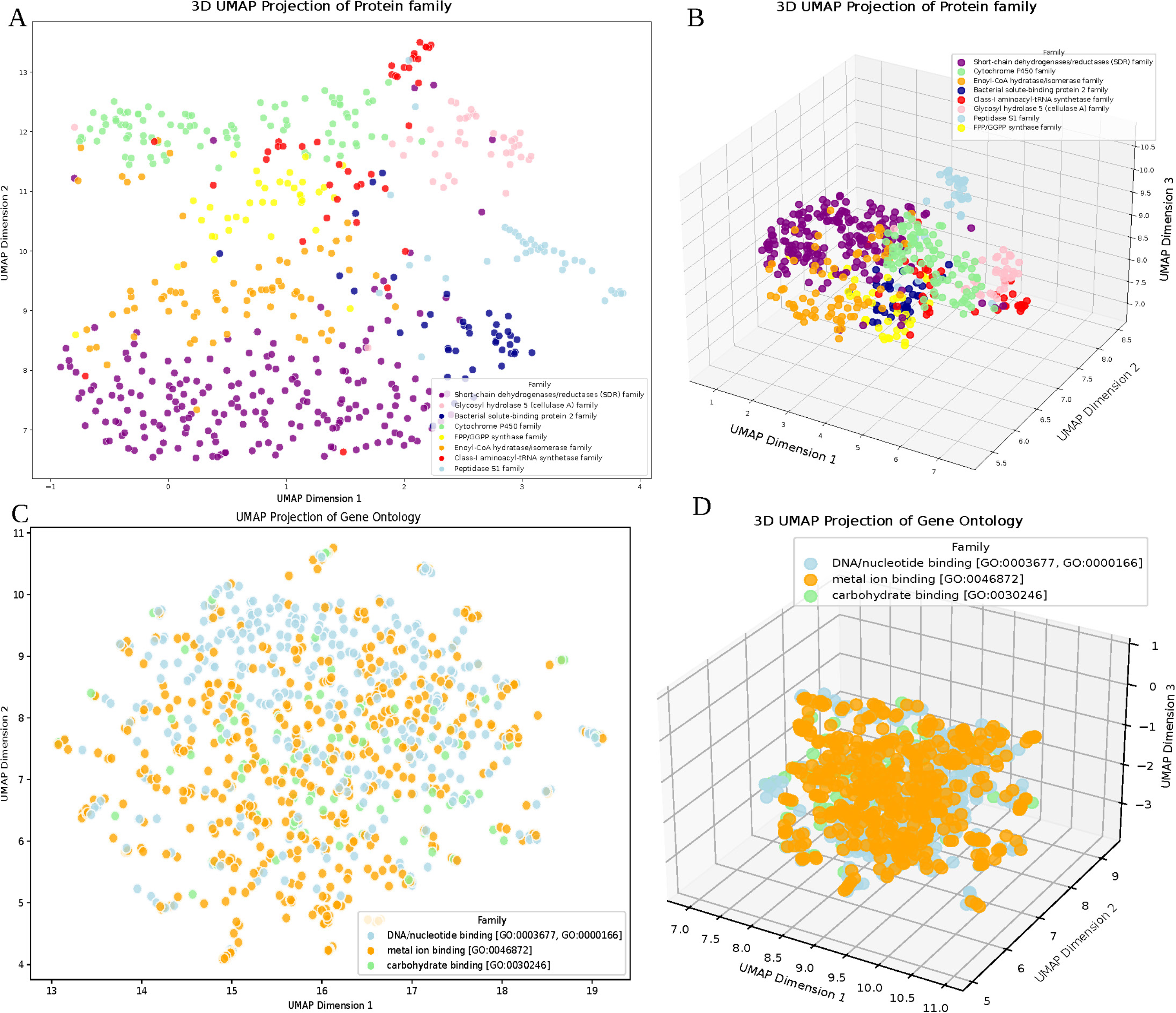
Pipeline of the InteracTor Algorithm. (A) Extensive database of PDB files, where each chain is separated into an individual PDB file. These files are converted to MOL2 format. (B) Molecular representation with carbon atoms in gray, oxygen in red, and nitrogen in blue. The following are calculated: Atom-atom distances (d) and van der Waals radii (r); Bond angles (a) and geometric centers (g); Interaction and atom types. (C) The following features are calculated: Interaction features (**Table 1**); Mono-, di-, and tripeptide composition; Side chain physicochemical properties. (D) The resulting data are saved in table format, which can be used later for feature selection, dimensionality reduction, clustering, visualization, and machine learning.

**Table 1:**
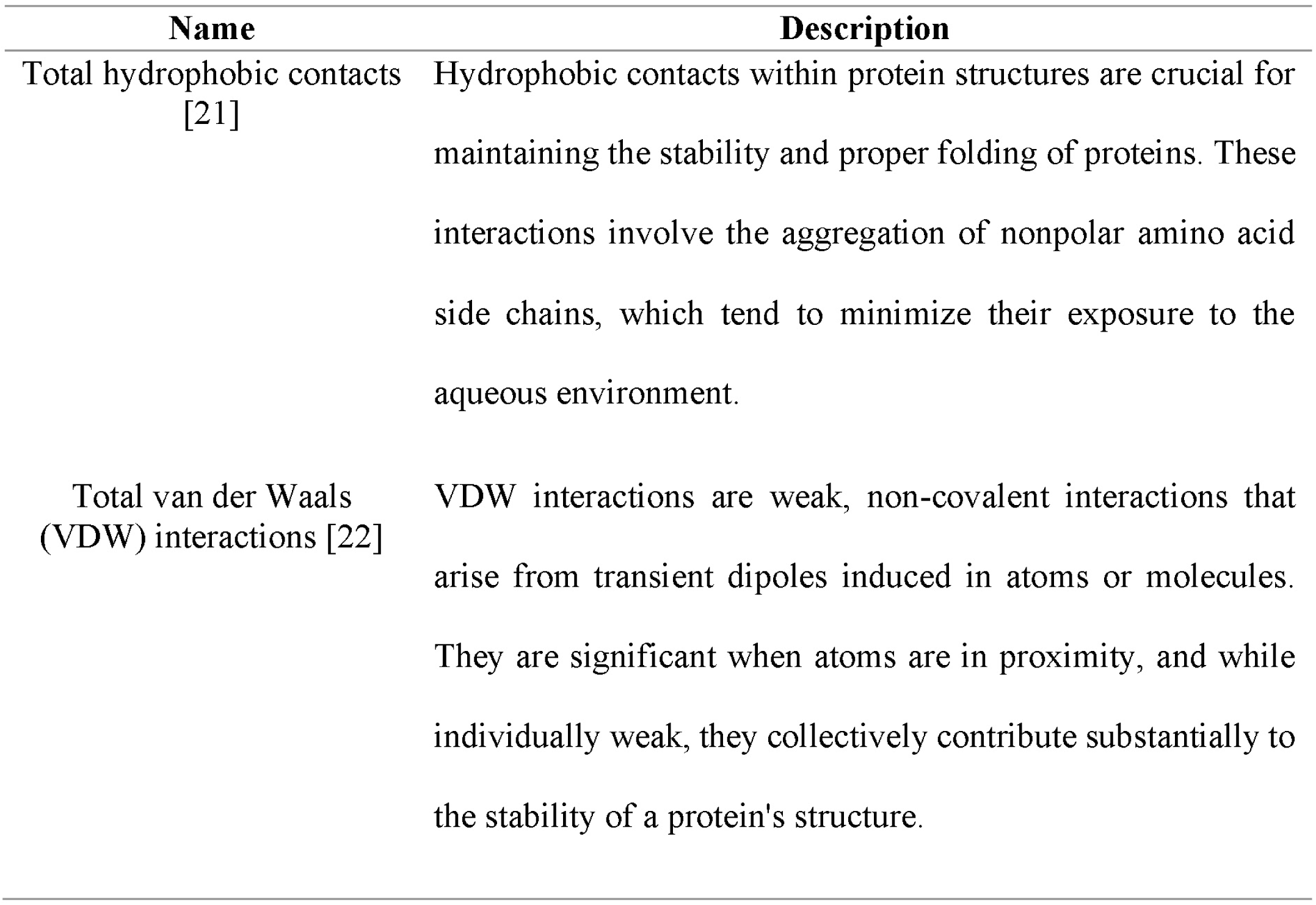
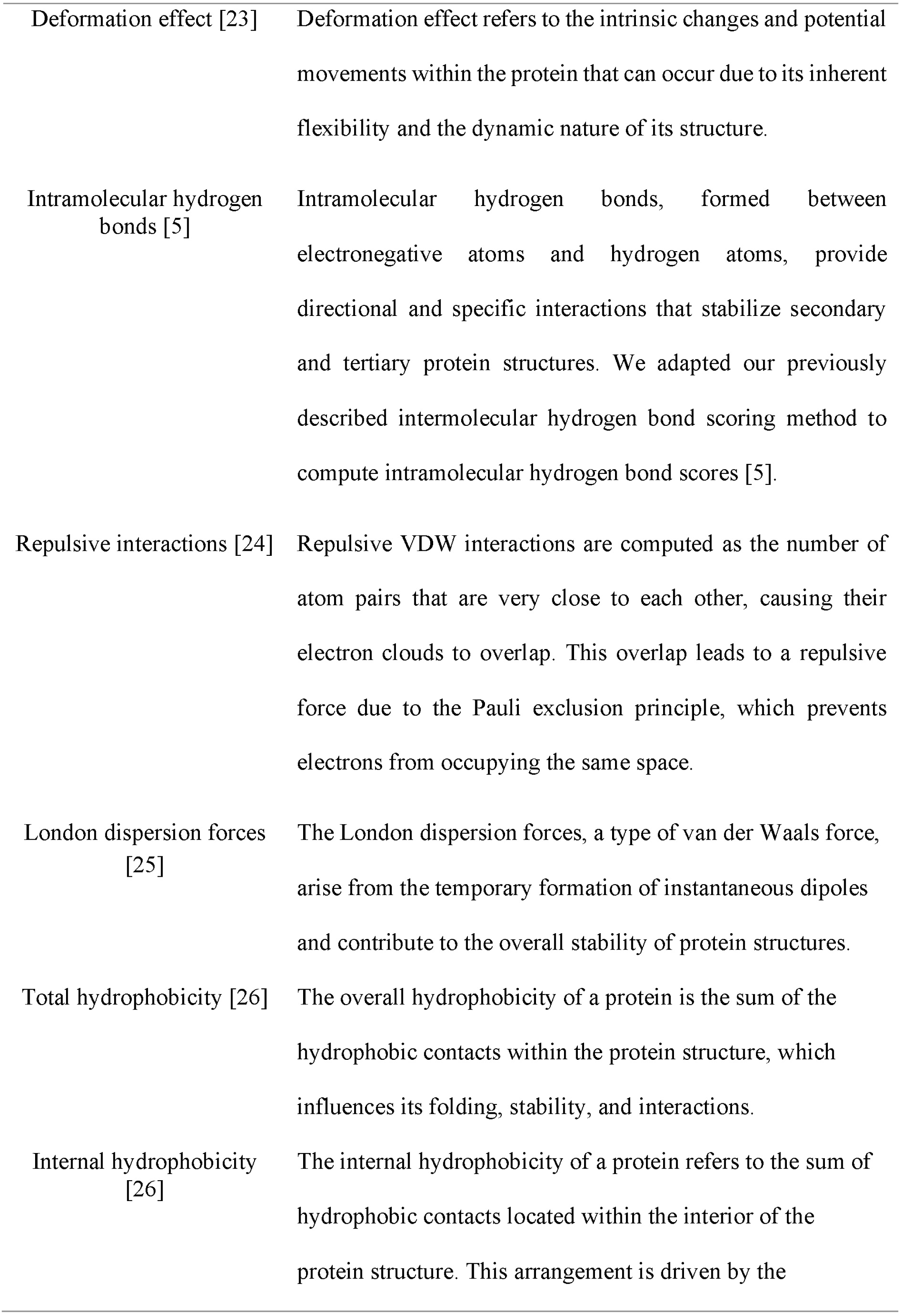
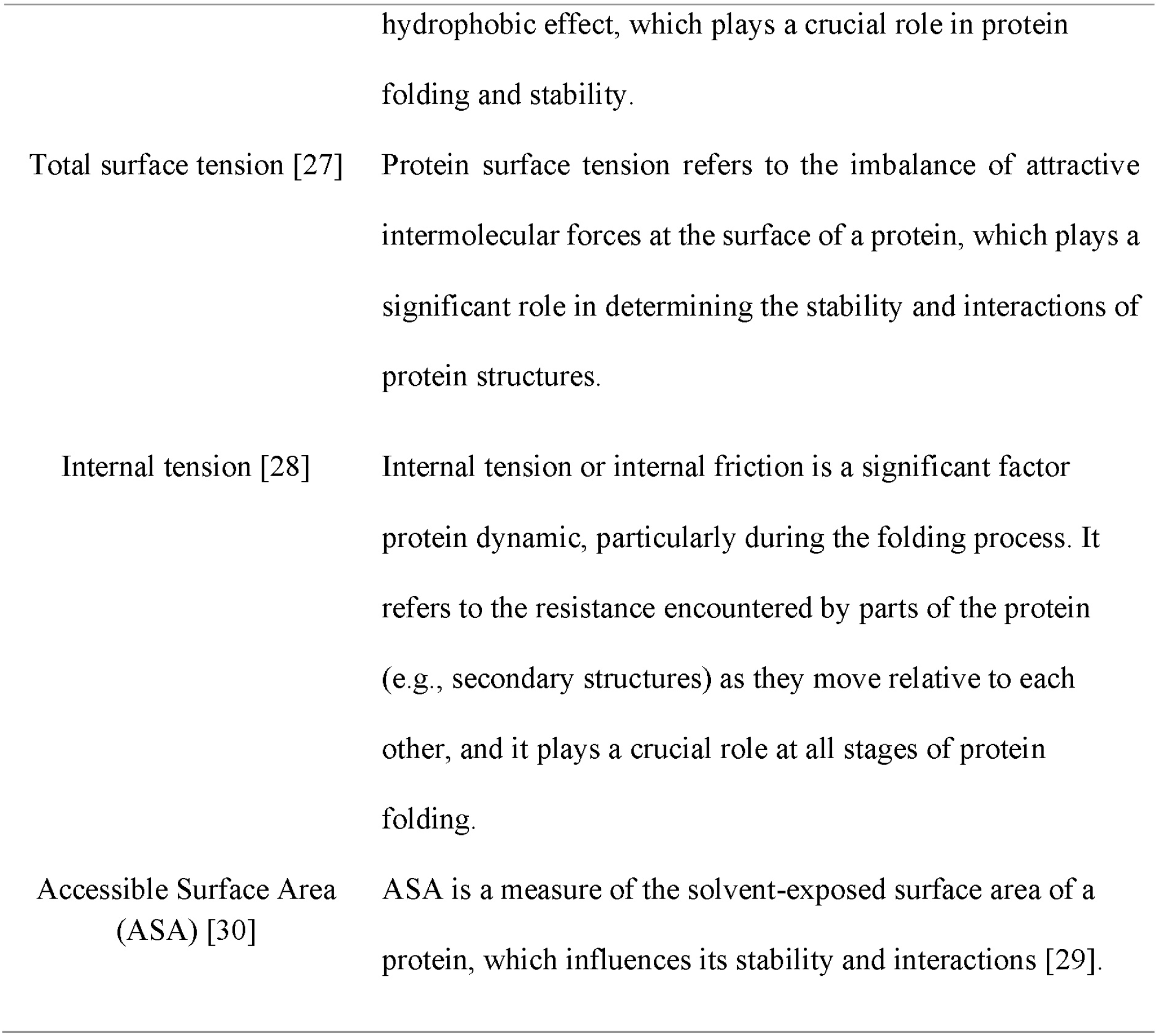
Protein structural features based interatomic interactions and protein physicochemical properties.

| Name | Description |
| --- | --- |
| Total hydrophobic contacts [21] | Hydrophobic contacts within protein structures are crucial for maintaining the stability and proper folding of proteins. These interactions involve the aggregation of nonpolar amino acid side chains, which tend to minimize their exposure to the aqueous environment. |
| Total van der Waals (VDW) interactions [22] | VDW interactions are weak, non-covalent interactions that arise from transient dipoles induced in atoms or molecules. They are significant when atoms are in proximity, and while individually weak, they collectively contribute substantially to the stability of a protein's structure. |
| Deformation effect [23] | Deformation effect refers to the intrinsic changes and potential movements within the protein that can occur due to its inherent flexibility and the dynamic nature of its structure. |
| Intramolecular hydrogen bonds [5] | Intramolecular hydrogen bonds, formed between electronegative atoms and hydrogen atoms, provide directional and specific interactions that stabilize secondary and tertiary protein structures. We adapted our previously described intermolecular hydrogen bond scoring method to compute intramolecular hydrogen bond scores [5]. |
| Repulsive interactions [24] | Repulsive VDW interactions are computed as the number of atom pairs that are very close to each other, causing their electron clouds to overlap. This overlap leads to a repulsive force due to the Pauli exclusion principle, which prevents electrons from occupying the same space. |
| London dispersion forces [25] | The London dispersion forces, a type of van der Waals force, arise from the temporary formation of instantaneous dipoles and contribute to the overall stability of protein structures. |
| Total hydrophobicity [26] | The overall hydrophobicity of a protein is the sum of the hydrophobic contacts within the protein structure, which influences its folding, stability, and interactions. |
| Internal hydrophobicity [26] | The internal hydrophobicity of a protein refers to the sum of hydrophobic contacts located within the interior of the protein structure. This arrangement is driven by the |
|  | hydrophobic effect, which plays a crucial role in protein folding and stability. |
| Total surface tension [27] | Protein surface tension refers to the imbalance of attractive intermolecular forces at the surface of a protein, which plays a significant role in determining the stability and interactions of protein structures. |
| Internal tension [28] | Internal tension or internal friction is a significant factor protein dynamic, particularly during the folding process. It refers to the resistance encountered by parts of the protein (e.g., secondary structures) as they move relative to each other, and it plays a crucial role at all stages of protein folding. |
| Accessible Surface Area (ASA) [30] | ASA is a measure of the solvent-exposed surface area of a protein, which influences its stability and interactions [29]. |

*<u>Step 1</u>.* Extract atom, residue, and sequence information from PDB file (**Figure 1A**): This step involves parsing the Protein Data Bank (PDB) file to obtain the atomic types, 3D coordinates, and the amino acid sequence of the protein.

*<u>Step 2</u>.* Extract additional information from MOL2 file (**Figure 1A**): In this step, we extract additional information from the protein’s atoms, including covalent bond mapping and types (e.g., single, double, triple bonds, aromatic rings, etc.) and sp^2^ hybridization.

*<u>Step 3.</u>* Calculate atom-atom distances (**Figure 1B**): In this step, the algorithm calculates the distances between pairs of atoms not covalently bound within the protein structure, which is used to identify potential interatomic interactions within the protein structure. A distance threshold, as described by Dias et al., [5] is applied to determine whether atoms are interacting. The minimum allowed distance between each pair of atoms is the sum of their van der Waals (VDW) radii minus 0.7, and the maximum allowed distance is the sum of their radii plus 0.7.

*<u>Step 4</u>.* Calculate geometric centers (**Figure 1B**): Additional properties are extracted from atoms selected as potentially participating in non-covalent interactions (step 3) and covalently bound to multiple atoms (step 2). This includes the calculation of geometric centers between the selected atoms and their respective covalently bound atoms. These centers are utilized to calculate angles between hydrogen bond donor and acceptor atoms (step 5).

*<u>Step 5.</u>* Calculate angles (**Figure 1B**): Here, the algorithm computes the angles between atoms or groups of atoms in the protein structure, which is used to evaluate hydrogen bonding geometry.

*<u>Step 6</u>*. Compute interatomic interaction features and protein physicochemical properties described in **Table 1** and **Figure 1C**.

*<u>Step 7.</u>* Extract compositional features (**Figure 1C**): InteracTor calculates the frequencies of single amino acids (monopeptide), pairs of amino acids (dipeptide), and triplets of amino acids (tripeptide) in the protein sequence.

*<u>Step 8</u>*. Extract CPAASC frequencies (**Table 2** and **Figure 1C**): This step involves calculating the frequencies of various physicochemical properties associated with the side chains of the amino acids in the protein sequence.

**Table 2:**
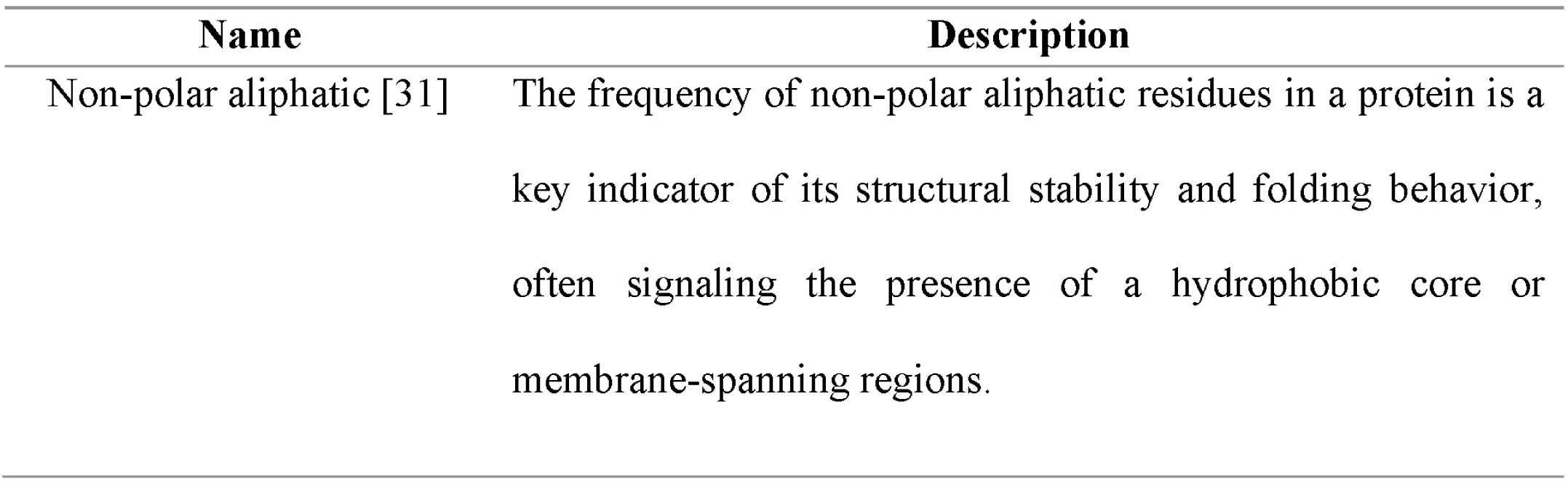
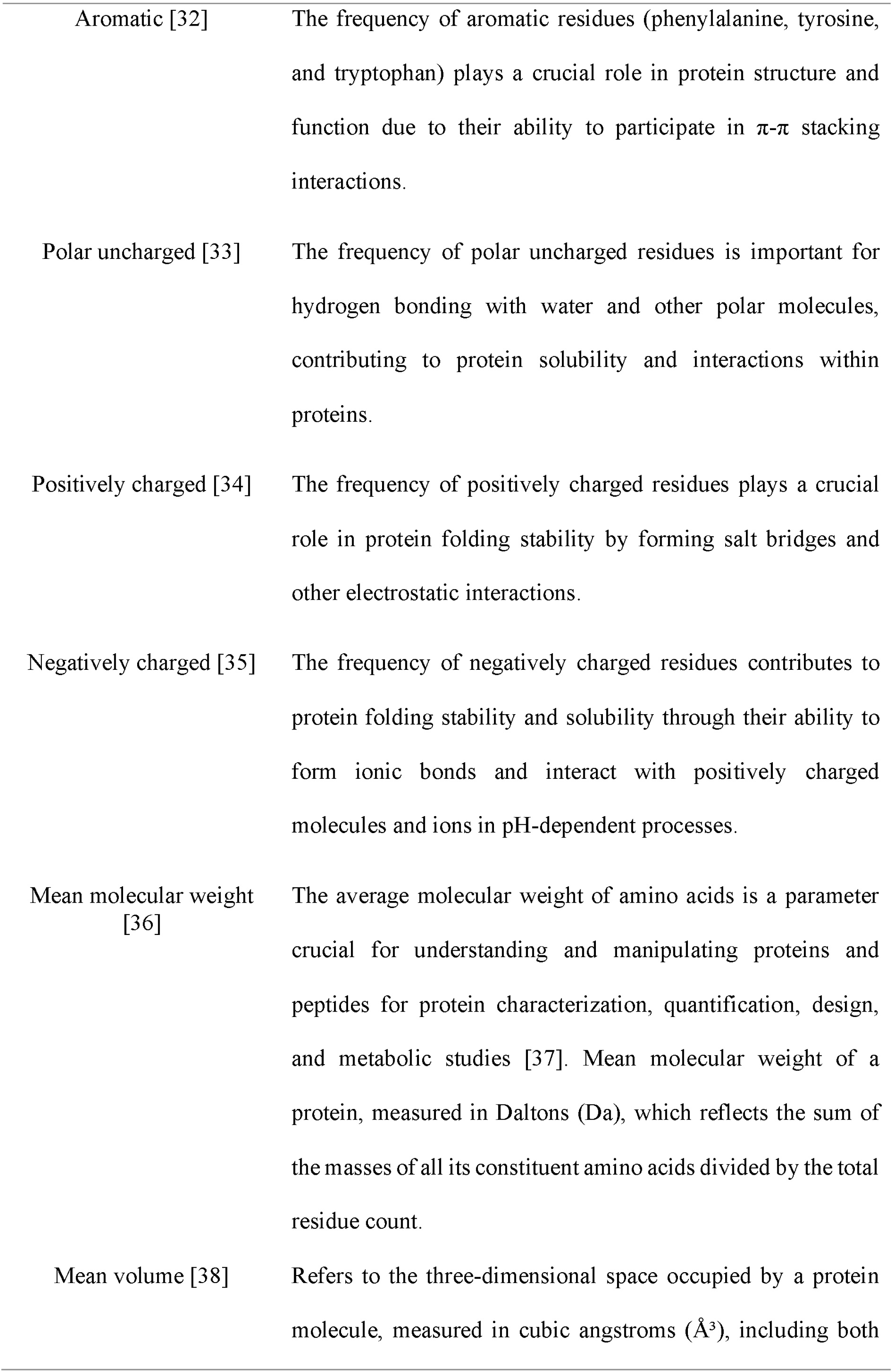
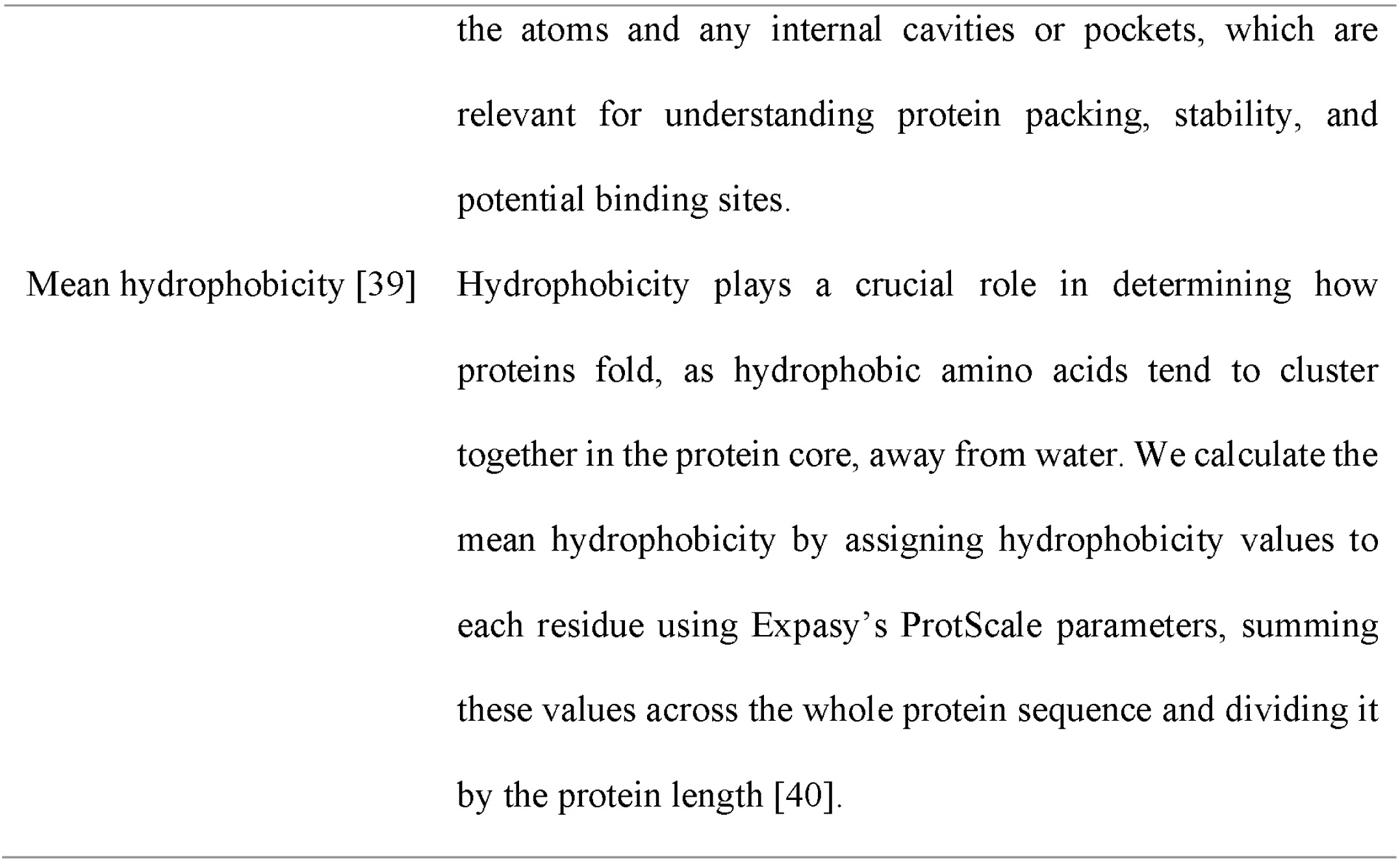
Protein physicochemical features based on chemical properties of amino acid side chains (CPAASC).

*<u>Step 9.</u>* Write results and postprocessing (**Figure 1D**): The final step is to write the features computed in steps 6, 7 and 8 to an output file for further analysis or use in other applications. The toolkit also includes example scripts for downstream analyses such as feature selection via MI scoring, dimensionality reduction via PCA, t-SNE, and UMAP, hierarchical clustering and visualization using heatmaps, and training machine learning algorithms.

### Implementation of protein structural features

We implemented a toolkit for the extraction and encoding of protein structural features based on physico-chemical properties, amino acids composition, and interatomic interactions. By building upon our previous methods for predicting protein-ligand binding affinity [5], we modified the algorithm to analyze residue-residue interactions within protein structures (**Table 1**). We also included the calculation of CPAASC frequencies [16] (**Table 2**), monopeptide [18], dipeptide, and tripeptide composition [17] (**Table 3**). In addition to the 20 classic amino acids, our algorithm supports rare residues found in distinct biological systems, including selenocysteine, pyrrolysine, and N-methylvaline. These amino acids, although rare, play important roles in specific biological processes [19,20].

**Table 3:**
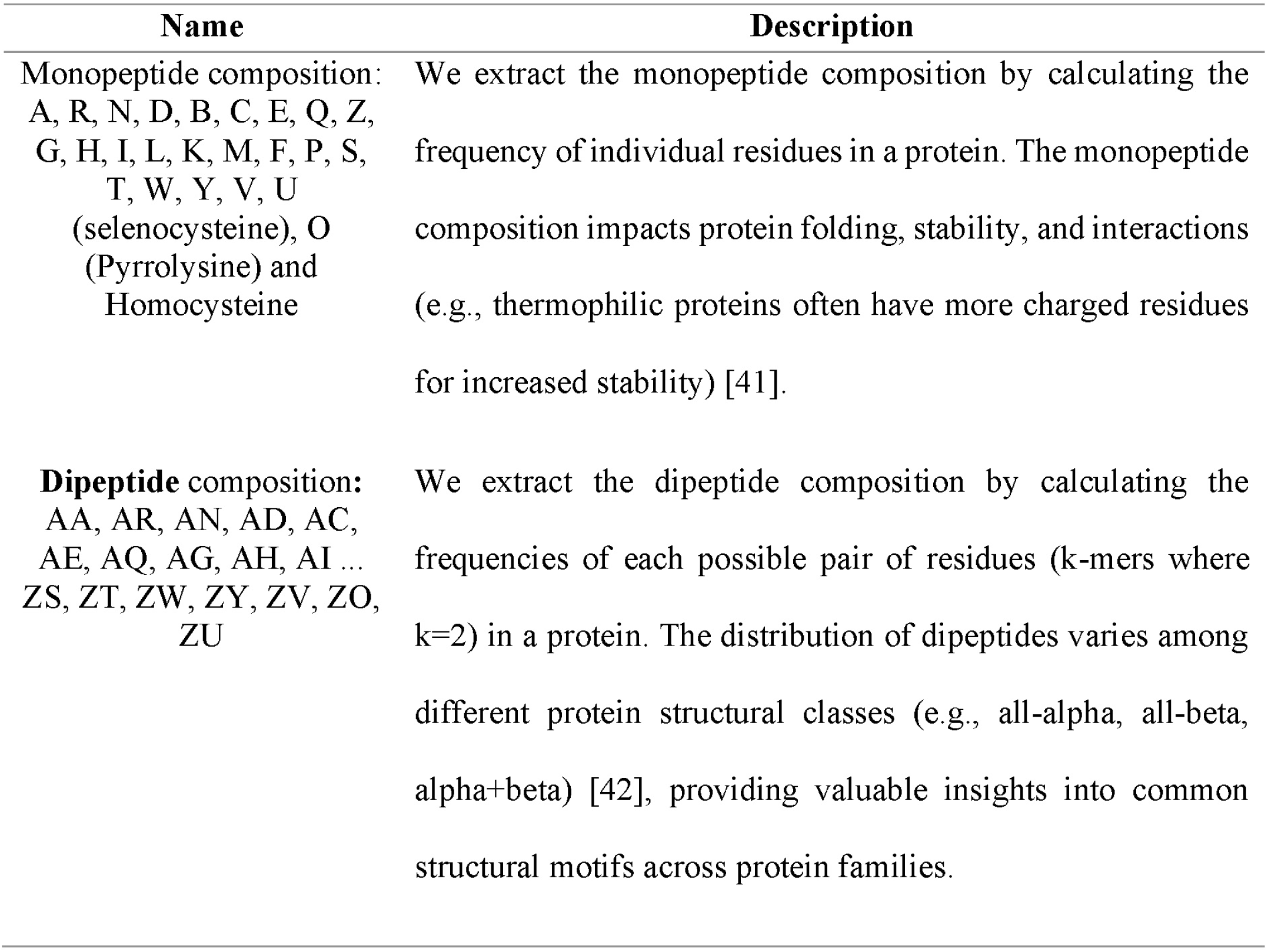
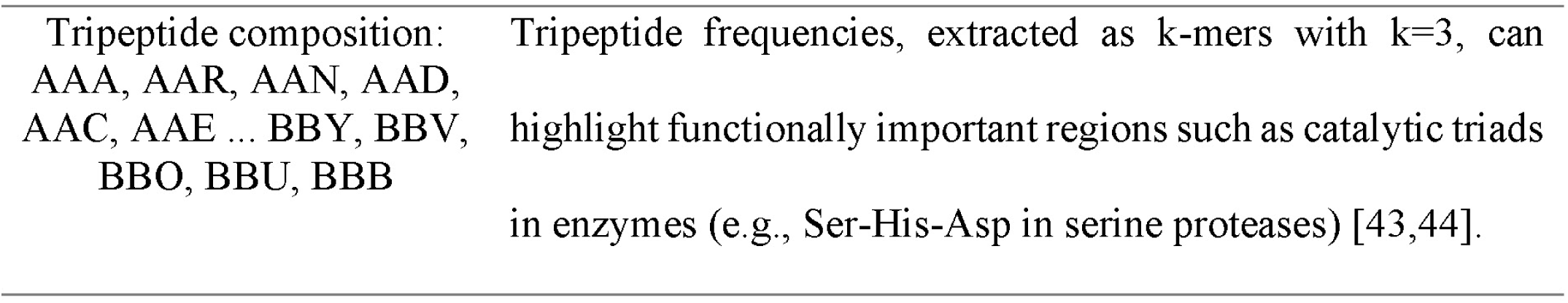
Protein sequence composition by k-mers frequencies features.

| Name | Description |
| --- | --- |
| **Monopeptide composition:**<br>A, R, N, D, B, C, E, Q, Z,<br>G, H, I, L, K, M, F, P, S,<br>T, W, Y, V, U<br>(selenocysteine), O<br>(Pyrrolysine) and<br>Homocysteine | We extract the monopeptide composition by calculating the frequency of individual residues in a protein. The monopeptide composition impacts protein folding, stability, and interactions (e.g., thermophilic proteins often have more charged residues for increased stability) [41]. |
| **Dipeptide composition:**<br>AA, AR, AN, AD, AC,<br>AE, AQ, AG, AH, AI ...<br>ZS, ZT, ZW, ZY, ZV, ZO,<br>ZU | We extract the dipeptide composition by calculating the frequencies of each possible pair of residues (k-mers where k=2) in a protein. The distribution of dipeptides varies among different protein structural classes (e.g., all-alpha, all-beta, alpha+beta) [42], providing valuable insights into common structural motifs across protein families. |
| Tripeptide composition:<br>AAA, AAR, AAN, AAD,<br>AAC, AAE ... BBY, BBV,<br>BBO, BBU, BBB | Tripeptide frequencies, extracted as k-mers with k=3, can<br>highlight functionally important regions such as catalytic triads<br>in enzymes (e.g., Ser-His-Asp in serine proteases) [43,44]. |

### Structural datasets and Preprocessing

We used 20,877 protein structures from the PDB-REDO database [45] eliminated redundant (100% sequence similarity) and small proteins (<50 residues) using Biopython and *in-house* scripts (see Code and Data Availability). PDB files were converted to MOL2 format using Open Babel [48]. We then applied the InteracTor algorithm to extract features from both PDB and MOL2 files.

### Data selected for demonstration of use cases

We utilized UniProt’s application programming interface (API) to extract Gene Ontology (GO) terms and protein family names based on the PDB accession number to facilitate analysis and demonstration of use cases [49,47]. We selected the most abundant protein families and GO terms in the dataset for further demonstration of use cases. Mapping scripts are available in GitHub as well (see Code and Data Availability).

### Feature selection

We applied MI scoring to quantify the relevance of each feature in distinguishing between different protein families and GO categories [50]. We ranked the features by their respective MI scores and selected the top 100 most informative features for further analyses.

### Analysis of protein structural families

We applied PCA to reduce the dimensionality of the data generated by our toolkit and evaluated how the primary components capture the variability among protein families and GO terms [51]. In addition to PCA, we also applied t-SNE [52,53,54] and UMAP [55] to reduce dimensionality while preserving nonlinear relationships and local structures in data. We used violin plots to visualize the distribution and density of the extracted features across different protein families and GO terms, providing insights into the distribution and central tendencies of the data [52].

We performed hierarchical clustering using Pearson correlation as distance metric and the complete linkage method for cluster formation [52]. This approach allowed us to identify and visualize the hierarchical relationships between protein families based on similarities among the features extracted by our toolkit. We then visualized the dimensionality reduction and clustering results with FreeViz [53,54]. For data analysis and visualization, the Python Data Mining Toolbox package was utilized [52].

## Results

InteracTor computes 11 different interatomic interaction features and 8 distinct CPAASC that are key for characterizing structure and function of proteins (**Table 1** and **Table 2**). In addition to interaction and structural features, our toolkit also extracts classic protein sequence composition features such as k-mer frequencies (**Table 3**), resulting in a total of 18,296 multimodal features extracted (**Table 4**). This provides a comprehensive representation of protein structure, function, and sequence characteristics, enabling in-depth analysis of protein properties across various scales of molecular organization.

**Table 4:**
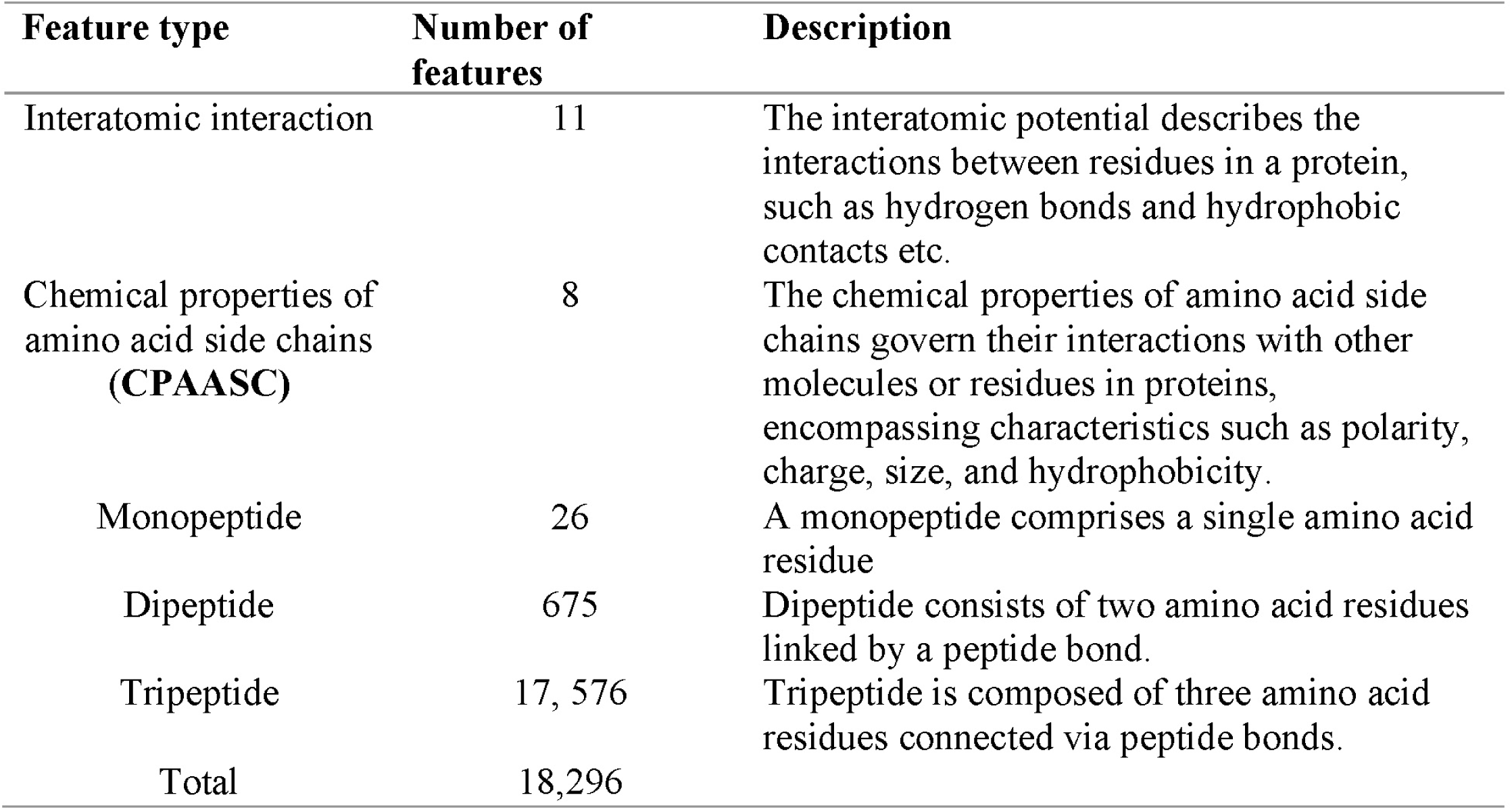
Overview of protein features extracted.

| Feature type | Number of features | Description |
| --- | --- | --- |
| Interatomic interaction | 11 | The interatomic potential describes the interactions between residues in a protein, such as hydrogen bonds and hydrophobic contacts etc. |
| Chemical properties of amino acid side chains (CPAASC) | 8 | The chemical properties of amino acid side chains govern their interactions with other molecules or residues in proteins, encompassing characteristics such as polarity, charge, size, and hydrophobicity. |
| Monopeptide | 26 | A monopeptide comprises a single amino acid residue |
| Dipeptide | 675 | Dipeptide consists of two amino acid residues linked by a peptide bond. |
| Tripeptide | 17, 576 | Tripeptide is composed of three amino acid residues connected via peptide bonds. |
| Total | 18,296 |  |

### InteracTor accurately characterizes the variability among distinct protein families

We conducted PCA across all 18,296 features extracted from 20,877 protein structures representing the most abundant protein families and GO terms in the PDB REDO database (**Supplementary Table 1**) to evaluate the overall utility of InteracTor in the characterization of variability among protein families and Gene Ontology (GO) terms. **Figure 2** shows the first two principal components, which captured 1.72% of the variance across protein families (**Figure 2B**), and 1.0% of the variance across GO terms (**Figure 2A**). Except for Peptidase S1 and the Glycosyl hydrolase 5 (cellulase A), the protein families exhibited well-defined clusters in the two-dimensional space (**Figure 2D**), whereas GO terms exhibited less separation overall (**Figure 2A**). This difference may be attributed to the inconsistent accuracy of GO annotations across non-model organisms [56]. Furthermore, features extracted from 3D structures could provide more accurate annotations compared to those based solely on sequence data, potentially improving the clustering and functional interpretation of protein families. Unlike the two-dimensional projections illustrated in Figure 2A and Figure 2B, the three-dimensional projections provide enhanced separation and clearer clustering of the data, allowing a distinct visualization of relationships among the protein structures. The first three principal components captured 2.40% of the variance across protein families (**Figure 2C, Figure S1A**), as well as 1.38% of the variance across GO terms (**Figure 2D, Figure S1B)**. Besides that, the 3D visualization improved the clustering and separation of Peptidase S1 and Glycosyl hydrolase 5 families, as well as the separation of GO categories, allowing for a more nuanced interpretation of functional relationships among groups (**Figure 2C**).

**Figure 2:**
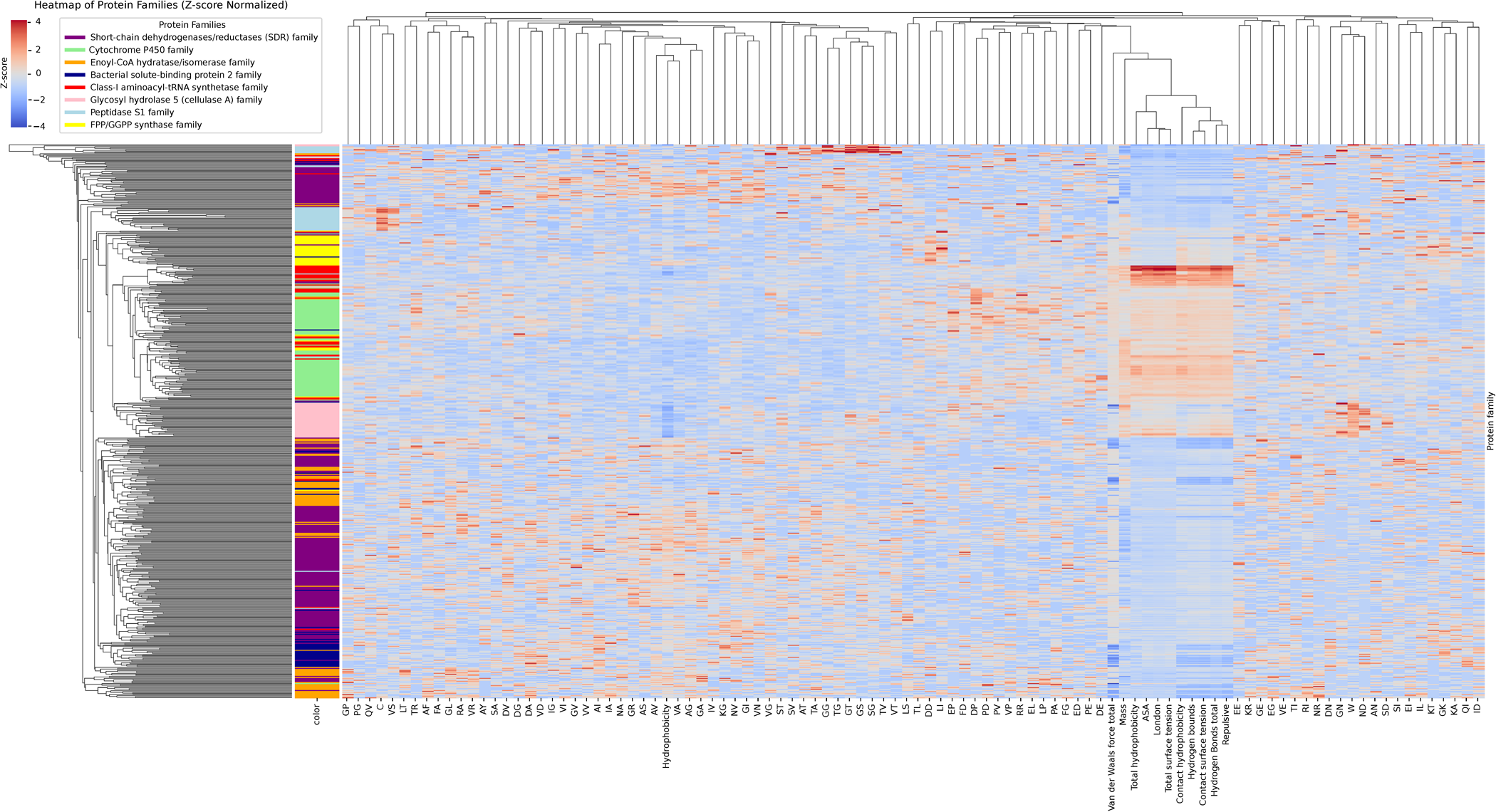
Analysis of Protein Families and Their Molecular Functions. A) Clustering of three representative groups based on molecular functions, emphasizing key functional categories and their interrelationships. B) Clustering of the most prominent protein family, showcasing distinct traits unique to that family. C) Three-dimensional PCA visualization of molecular functions, capturing the variance and complexity within different functional categories. D) Three-dimensional PCA representation of protein families, showing clear separations and relationships among them.

### Mutual Information scoring identifies key features across protein families and GOs

In the feature selection process, all features associated with the selected protein families were ranked using the MI score. The histogram of MI scores used for feature selection shows a bimodal distribution (**Figure 3A**). The primary mode, or large peak, corresponds to features with low MI scores (MI<0.2), reflecting background noise and less informative features. The remaining lower peaks (MI>=0.2) represent a subset of 354 features with informational content effectively distinguished from the background noise peak. Among the top 100 high MI scoring features are 9 interatomic interactions, 2 CPAASC features, and 89 sequence composition features (**Supplementary Table 3**). **Figure 3B** displays the distribution of MI scores across all features, highlighting the spread and density of these scores. Among the 12 most highly ranked features across protein families are hydrogen bonds (MI=0.775), total surface tension (MI=0.763), london dispersion forces (MI=0.758), repulsive interactions (MI=0.722), internal tension (MI=0.708), ASA (MI=0.694), hydrophobic contacts (MI=0.561), TG frequency (MI=0.562), internal hydrophobicity (MI=0.561), VN frequency (MI=0.556), total hydrophobicity (MI=0.539), and GG frequency (MI=0.509). For GO terms, the MI score analysis revealed values ranging from approximately 0.06 to 0.1, as shown in **Figure 3C**. While several peaks are well-defined, indicating key features, the MI scores are notably lower compared to those for protein families. **Figure 3D** presents a violin plot illustrating the distribution of MI scores, with a high density of values between 0.1 and 0.2. High MI scoring features include total surface tension (MI=0.202), london dispersion forces (MI=0.165), total hydrophobicity (MI=0.146), ASA (MI=0.142), EL frequence (MI=0.127), VDW interactions (MI=0.123), EG, NL, KT, GN, AN, (MI=0.113, 0.111, 0.109, 0.105, and 0.103, respectively) and repulsive interactions (MI=0.100). Despite the lower MI scores, these features demonstrate a significant (Wilcoxon test, p-value < 2.2^−16^) influence on the differentiation of protein family and GO categories, highlighting their relevance in the context of ligand/substrate interactions and functional protein family.

**Figure 3:**
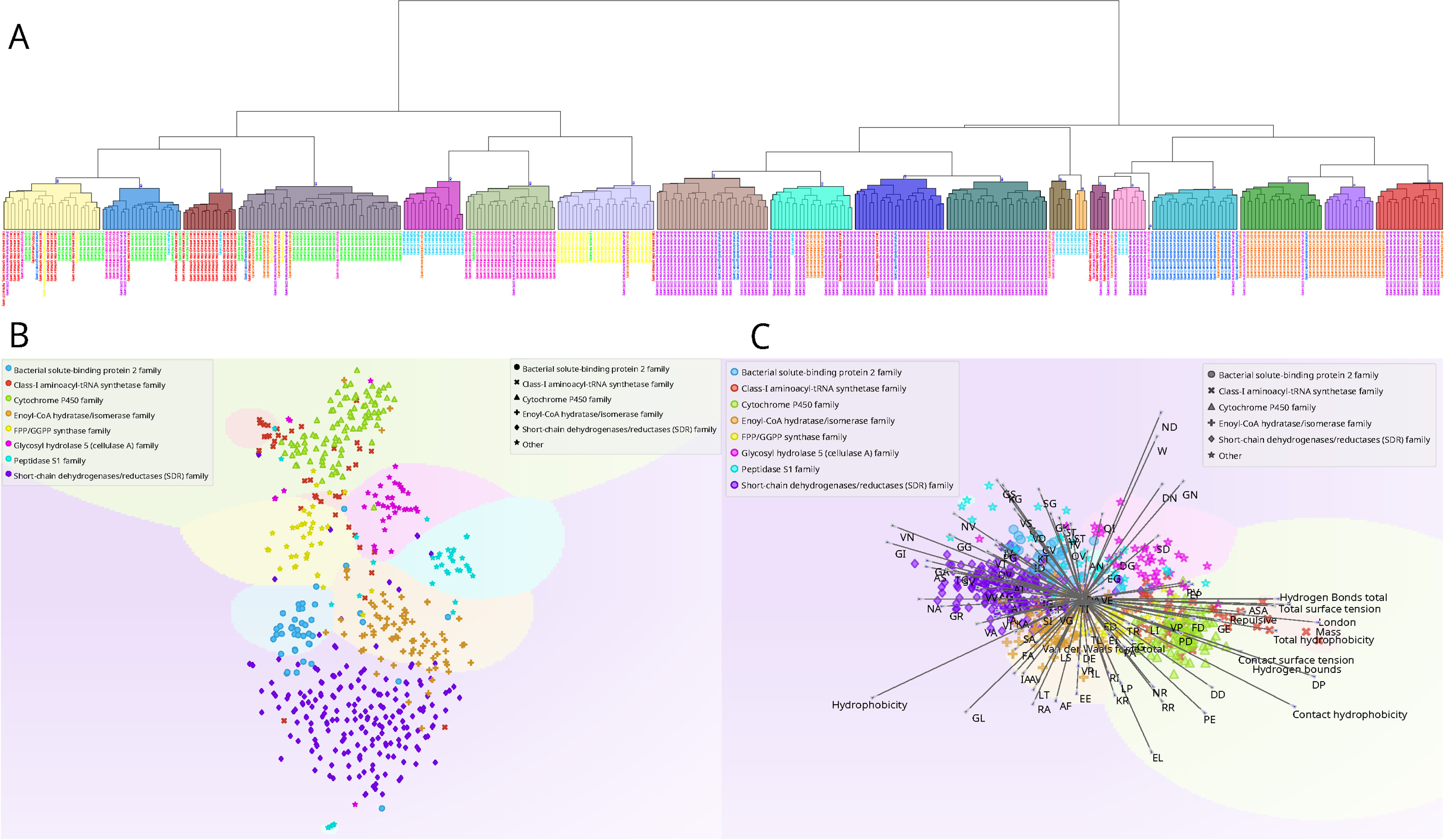
Analysis of Mutual Information (MI) Scores Across Protein Families and Molecular Functions. A) Frequency distribution of MI scores for each protein family B) Density distribution of MI scores within protein families, focusing on the top 12 ranked features that significantly contribute to family differentiation. C) Frequency distribution of MI scores for molecular functions. D) Density distribution of MI scores for molecular functions, emphasizing the top 12 ranked features.

### Selected features reveal unique and distinct distributions across protein families and GOs

**Figure 4A** shows consistent distribution of london dispersion forces and total surface tension features across multiple protein families, including Peptidase S1, Cytochrome P450, and Glycosyl Hydrolase 5. In **Figure 4B** and **4D**, the repulsive interactions and hydrogen bonds features show similar patterns among Bacterial Solute Binding Protein 2, Enoyl-CoA Hydratase/Isomerase, and Glycosyl Hydrolase families, displaying a bimodal distribution. Further analysis of the repulsive interactions and hydrogen bond features (**Figure 4B** and **D**, respectively) revealed similar distribution patterns across the analyzed families (Spearman correlation, r = 0.99, p = 0.968, Supplementary Figure S2). Bacterial Solute Binding Protein 2, Enoyl-CoA Hydratase/Isomerase, and Glycosyl Hydrolase families displayed bimodal distributions.

**Figure 4:**
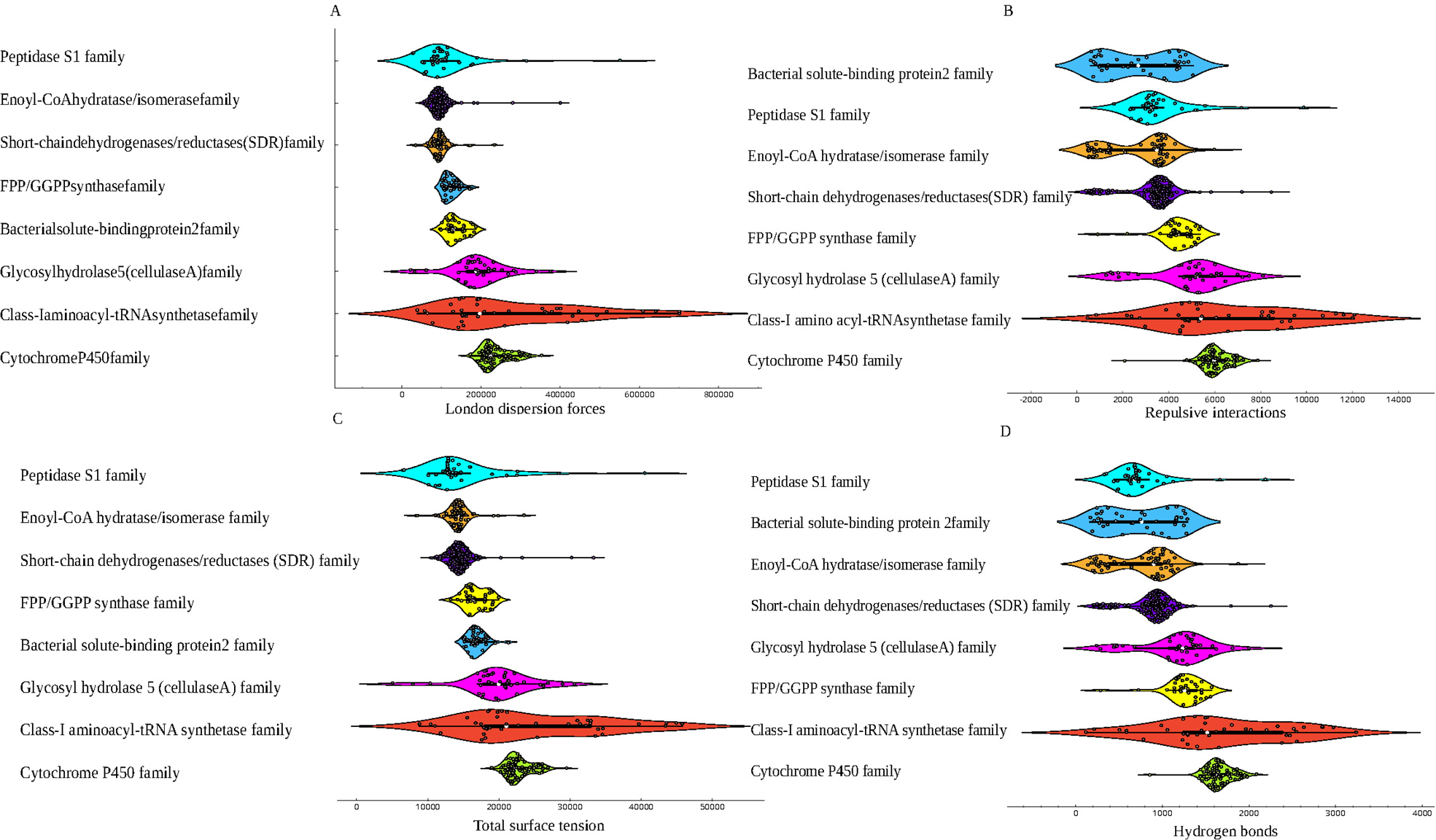
Distribution and Frequency of Key Features Across Protein Families. A) Distribution and frequency of the London dispersion feature among various protein families. B) Distribution and frequency of the repulsive interaction feature across different protein families. C) Distribution and frequency of the total surface tension feature for each protein family. D) Distribution and frequency of the total hydrogen bond count within protein families.

### Clustering of selected features reveals distinct patterns and relationships among protein families

Hierarchical clustering identified 19 distinct groups across eight families (**Figure 5A**), revealing a complex internal structure within the dataset. Complementary t-SNE clustering analysis (**Figure 5B**) detected 9 groups and not only corroborated these groupings but also unveiled finer details of inter-group relationships. While hierarchical clustering effectively segmented proteins with similar functions, t-SNE provided a more nuanced separation, highlighting occasional cross-family similarities, with some datapoints from different families appearing in unexpected clusters. Multivariate visualization (**Figure 5C**) further highlighted the role of interatomic interaction features, which were predominantly observed in the Cytochrome P450 and Glycosyl Hydrolase 5 (cellulase A) families (Wilcoxon p ≤ 0.00049, See **Supplementary Table 4**). In contrast, dipeptide features were more prevalent among other protein families, indicating a varied functional landscape shaped by distinct sequence composition patterns. (Wilcoxon p ≤ 0.03125, See **Supplementary Table 4**).

**Figure 5:**
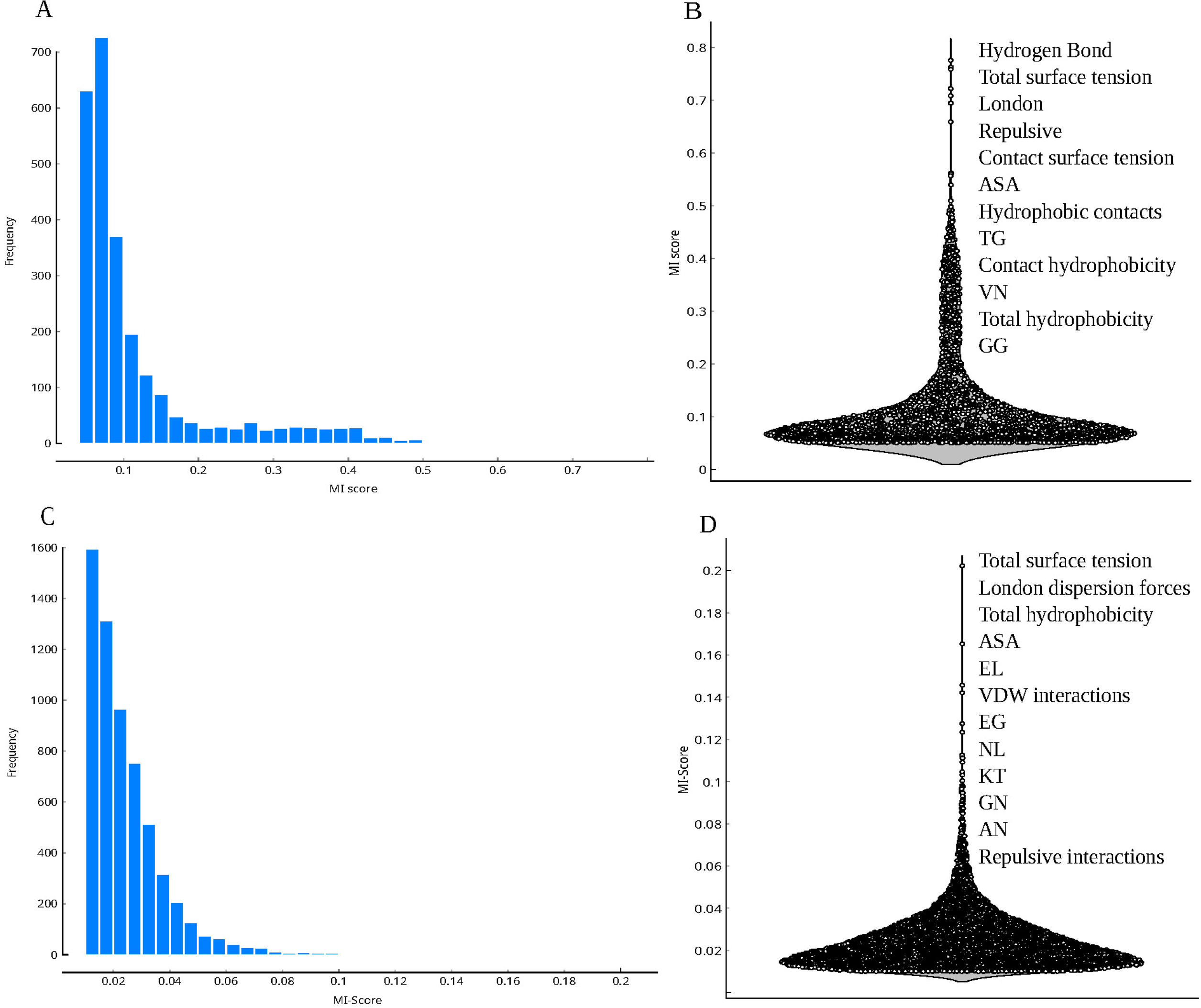
Advanced Clustering and Visualization of Protein Families. A) Hierarchical clustering of protein families, depicting relationships and groupings based on structural and functional similarities. B) t-SNE plot offering a refined visualization of inter-family relationships within a reduced-dimensional space. C) FreeViz plot highlighting the most significant features for each group, illustrating key attributes that differentiate protein families.

## Discussion

### A Novel Algorithm for Improved Characterization of Protein Structural Properties

Our protein feature extraction algorithm distinguishes itself by focusing on the tertiary structure of proteins, particularly on interatomic interactions, unlike other approaches that rely on primary and secondary structures derived from amino acid sequences [10, 57–58]. By analyzing three-dimensional relationships, the algorithm captures intramolecular interactions that underpin protein structure and function. Extracting features directly from the tertiary structure allows the algorithm to uncover additional patterns in the chemical properties of amino acids and their interactions, offering a comprehensive understanding of the structural and functional dynamics of proteins [59]. This enhanced perspective provides new insights into the complex mechanisms governing protein behavior, facilitating advancements in protein engineering and drug design [60]. The graphical representations (**Figures 2, 5, 6, and 7**) provide a comprehensive overview of protein characteristics as well, assisting in the visualization and interpretation of complex structural and physicochemical properties across protein families.

**Figure 6:**
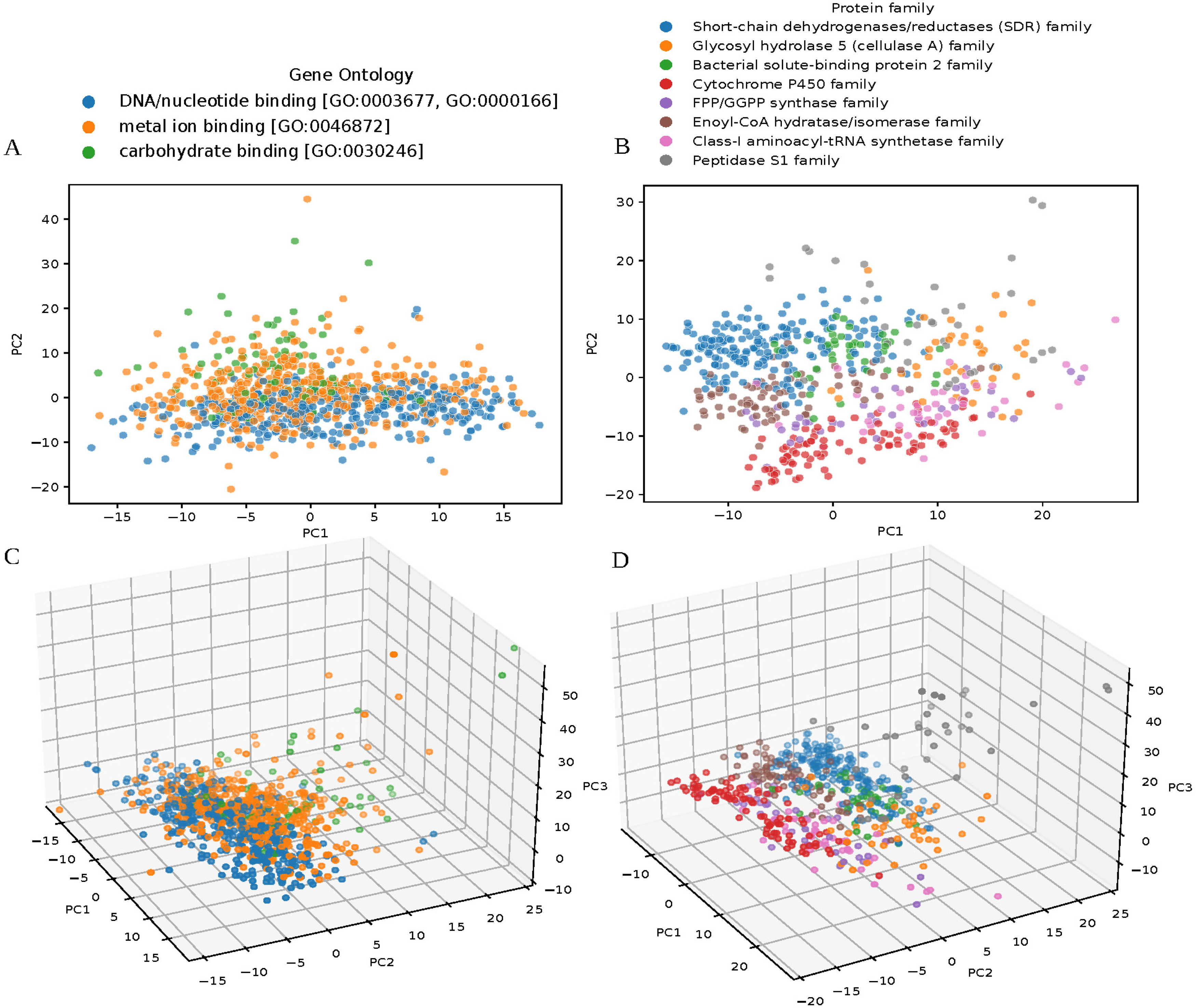
Heatmap of Protein Family and Feature Clustering. Heatmap displays the clustering of protein families and features, emphasizing interatomic interaction characteristics. Z-scores were applied to standardize the data, highlighting distinct clusters based on these pivotal interactions.

**Figure 7:**
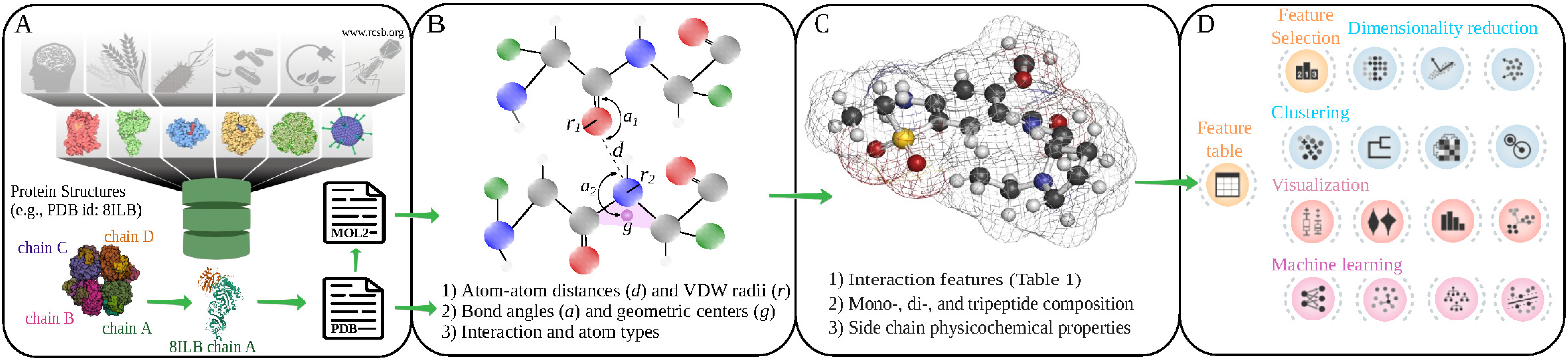
Two-Dimensional Projection of Protein Families. This UMAP projection visualizes the clustering of protein families, showcasing distinct groupings informed by structural and functional characteristics.

The feature selection process, guided by MI scores, was crucial in identifying the most effective features for distinguishing protein families, as shown in **Figure 3A** and **B**. Prominent peaks in MI scores indicate that features such as hydrogen bonds, total surface tension, and contact hydrophobicity are particularly influential in differentiating protein families, underscoring their critical role in our analysis. This prioritization of key features enhances our ability to effectively distinguish between various protein structural and functional groups. The identification of 9 out of 11 interatomic interactions features among the top 12 features with the highest MI scores underscores their critical role in differentiating protein families. This finding defines the importance of specific molecular interactions, such as hydrogen bonds and hydrophobic contacts, in defining protein structure and function, providing valuable insights into the fundamental principles governing protein family diversity.

**Figure 6** revealed clear definitions of specific clusters, highlighting the effectiveness of the approach in identifying grouping patterns among protein families. The groupings not only reflected complex interatomic relationships between different features but also demonstrated the formation of distinct clusters within dipeptide feature groups. The Peptidase S1 family, Glycosyl hydrolase 5 (cellulase A) family and Short-chain dehydrogenases/reductases (SDR) family demonstrated clustering of dipeptide feature groups. This result confirms the method’s ability to highlight relevant functional patterns and distinguish between different protein categories based on the observed interatomic interactions.

A bimodal distribution of hydrogen bonding in a protein family may suggest the presence of two distinct subtypes of structural or functional groups within that family (**Figure 4D**). This variability might arise from differences in the protein’s environment or structure, such as variations in hydrophobicity or the presence of neighboring groups that influence hydrogen bond strength and stability. For instance, hydrogen bonds can be stronger in hydrophobic environments due to lower dielectric constants, and their contributions to stability can vary significantly depending on their distance, geometry, and surrounding interactions. Additionally, a bimodal distribution might suggest the presence of a miss annotation or two subfamilies within the protein family, each characterized by distinct hydrogen bonding patterns that could be related to specific structural or functional adaptations (**Figure 4**).

Visualization of the two-dimensional projection generated via UMAP analysis (**Figure 7**) separated key protein families, including Short-chain dehydrogenases/reductases (SDR), Cytochrome P450, Enoyl-CoA hydratase/isomerase, Bacterial solute-binding protein 2, Class-I aminoacyl-tRNA synthetase, Glycosyl hydrolase 5 (cellulase A), Peptidase S1, and FPP/GGPP synthase. Each family formed distinct clusters, indicating that UMAP effectively distinguished these groups based on the protein features extracted by InteracTor. This clear separation underscores the method’s ability to reveal meaningful patterns and relationships within the dataset, offering valuable insights into the functional dynamics of the protein families.

### Case Study on Multi-dimensional Clustering Reveals Complex Structure and Functional Diversity in Protein Families

Hierarchical clustering effectively delineated differences among protein families such as short-chain dehydrogenases/reductases (SDR), Cytochrome P450, Enoyl-CoA hydratase/isomerase, Bacterial solute-binding protein 2, Class-I aminoacyl-tRNA synthetase, Glycosyl hydrolase 5 (cellulase A), Peptidase S1, and FPP/GGPP synthase. Complementary analysis with t-SNE allowed the detection of subtle similarities and differences among the protein families and offering deeper insights into their functional connections (**Figure 5**, See supplementary **Table 4**). Multivariate visualization allowed dynamic adjustments to identify unique aspects of protein families, including specific features contributing to the clustering process, yielding valuable insights into the functional diversity within and between protein families (**Figure 5, 6, 7**). Additionally, UMAP captured both local and global structures in a reduced-dimensional space, allowing a comprehensive understanding of the clustering and distribution of protein families, highlighting both shared and similar structural characteristics. This integrative approach led to a deeper understanding of the complex relationships and functional diversity inherent across protein families, revealing details on the interplay of intermolecular interactions, physicochemical properties, and composition of proteins. This holistic methodology not only enhances our comprehension of protein family dynamics but also establishes a robust framework for future research. The extracted features have the potential to directly impact the performance of machine learning and deep learning models by offering a new feature engineering tool. By providing new insights on protein structural and functional properties, our approach supports advancements in structural biology, synthetic biology and in the characterization proteins of unknown function (PUFs) [61, 62, 63, 64]. Additionally, these features can contribute to emerging fields such as quantum biology [64, 65, 66, 67, 68], facilitating a deeper understanding of biological roles and interactions across diverse protein families.

## Supporting information

Supplementary Table 1: Distribution and quantification of selected protein families

Supplementary Table 2: Distribution and quantification of selected gene ontology

Figure S1. Principal component of protein families and GO terms. (A) PCA plot illustrating the variance captured by the first three principal compone

Figure S2. Analysis of correlation between repulsive interactions and hydrogen bonds. (A) Correlation analysis based on the features of repulsive inte

Supplemental Table 3. MI Scores for protein families and GO Terms.

Supplemental Table 4. Wilcoxon p-values for feature pairwise comparisons between protein families.

## Key Points

- This paper provides a comprehensive toolkit for extracting and encoding sequence, physicochemical, and structural features from proteins.
- The proposed toolkit introduces a novel algorithm for extracting interatomic interaction features from protein structures.
- This work also provides detailed use cases, demonstrating how our toolkit can enhance protein family characterization of distinct protein families by leveraging several types of clustering algorithms.
- Our comprehensive feature extraction and analytical methods revealed new insights into protein structure-function relationships.
- The new toolkit is available as an open-source software on GitHub, promoting accessibility, reproducibility and collaboration in research.

## Acknowledgements

The authors gratefully acknowledge UF Research Computing for providing computational resources and support that have contributed to the research reported in this publication (http://www.rc.ufl.edu) and the financial support provided by Launching Innovative Faculty Teams in AI (LIFT AI) program.

## Author Contributions

R.D. designed and supervised the study and contributed to manuscript writing. M.F.R.R and

M.K. contributed to manuscript writing. J.C.F.S. developed the algorithm, built the computational tools, performed the analysis, and wrote the manuscript. L.S and N.S. were responsible for the data preprocessing and contributed to manuscript writing. All authors read and approved the final manuscript.

## Code and Data Availability

InteracTor is provided as an open-source python in GitHub: https://github.com/Dias-Lab/InteracTor

## Supplementary Figures

**Figure S1. Principal component of protein families and GO terms.**

(A) PCA plot illustrating the variance captured by the first three principal components across different protein families. (B) PCA plot depicting the variance explained by the first three principal components across Gene Ontology (GO) terms.

**Figure S2. Analysis of correlation between repulsive interactions and hydrogen bonds.**

(A) Correlation analysis based on the features of repulsive interactions and hydrogen bonds.

(B) Average values of hydrogen bonds across different conditions. (C) Average values of repulsive interactions across different conditions.

## Supplementary Tables

**Supplementary Table 1:**
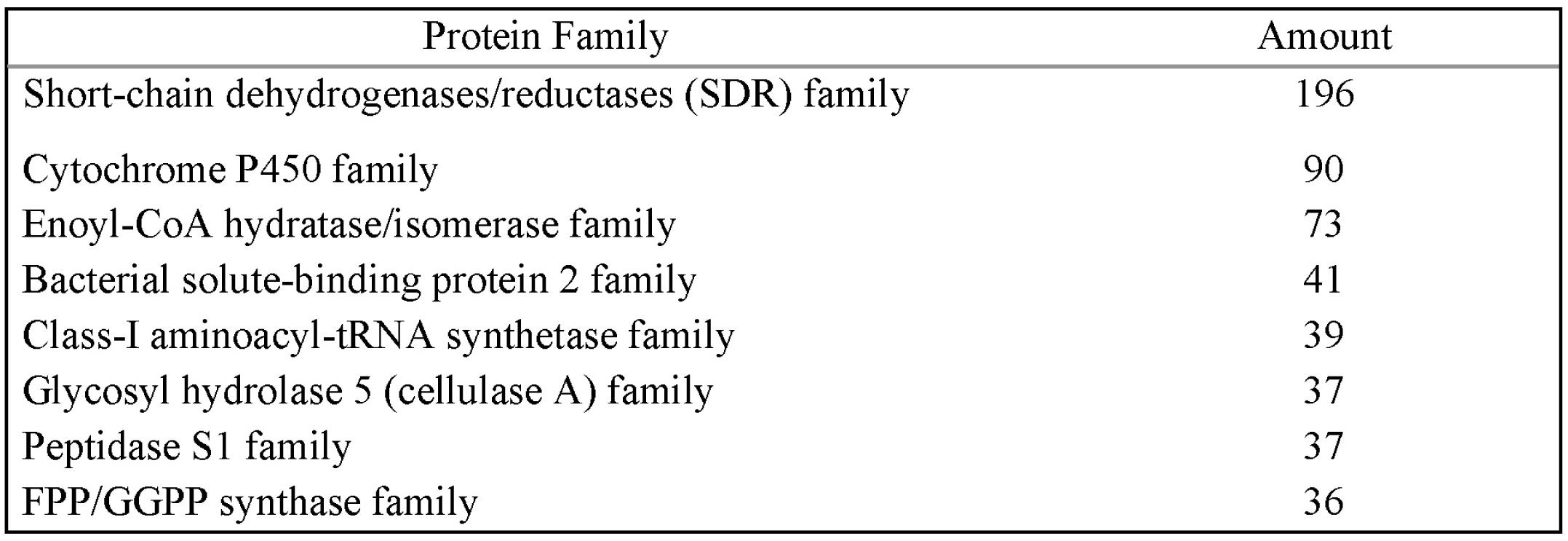
Distribution and quantification of selected protein families.

**Supplementary Table 2:**
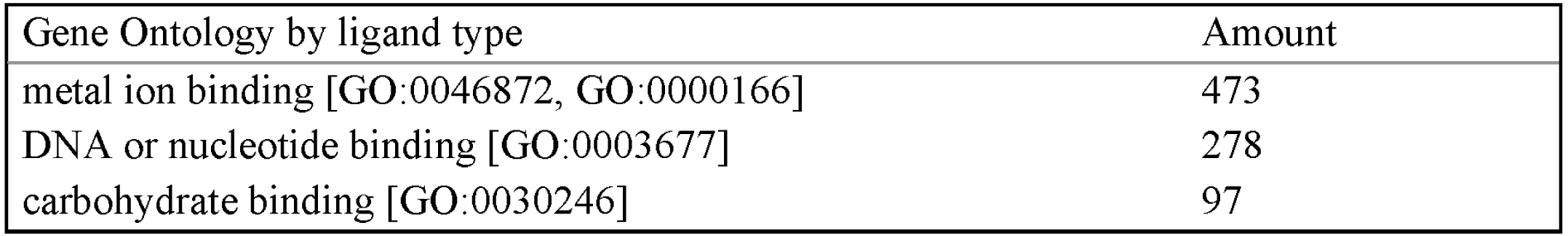
Distribution and quantification of selected gene ontology.

| Gene Ontology by ligand type | Amount |
| --- | --- |
| metal ion binding [GO:0046872, GO:0000166] | 473 |
| DNA or nucleotide binding [GO:0003677] | 278 |
| carbohydrate binding [GO:0030246] | 97 |

**Supplemental Table 3. MI Scores for protein families and GO Terms.**

**Supplemental Table 4. Wilcoxon p-values for feature pairwise comparisons between protein families.**

