## Supplementary Table 1: Distribution and quantification of selected protein families for "InteracTor: A new integrative feature extraction toolkit for improved characterization of protein structural properties"

| Protein Family | Amount |
|---|---|
| Short-chain dehydrogenases/reductases (SDR) family | 196 |
| Cytochrome P450 family | 90 |
| Enoyl-CoA hydratase/isomerase family | 73 |
| Bacterial solute-binding protein 2 family | 41 |
| Class-I aminoacyl-tRNA synthetase family | 39 |
| Glycosyl hydrolase 5 (cellulase A) family | 37 |
| Peptidase S1 family | 37 |
| FPP/GGPP synthase family | 36 |
