## Supplementary figures and images for "InteracTor: A new integrative feature extraction toolkit for improved characterization of protein structural properties"

### Figure S1. Principal component of protein families and GO terms. (A) PCA plot illustrating the variance captured by the first three principal compone

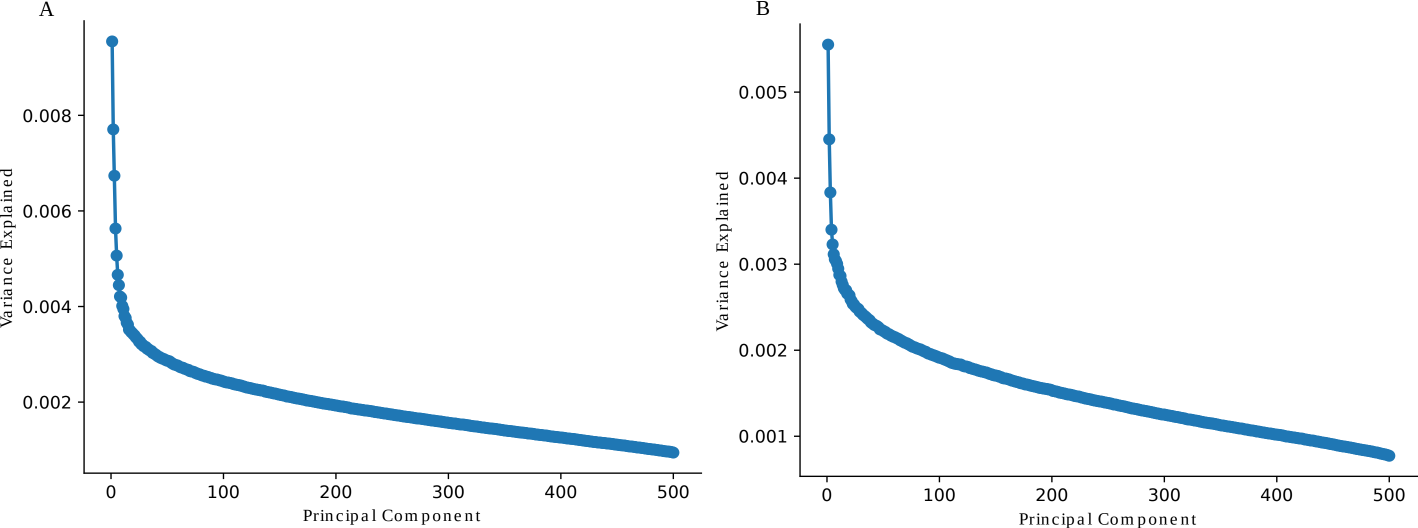

### Figure S2. Analysis of correlation between repulsive interactions and hydrogen bonds. (A) Correlation analysis based on the features of repulsive inte

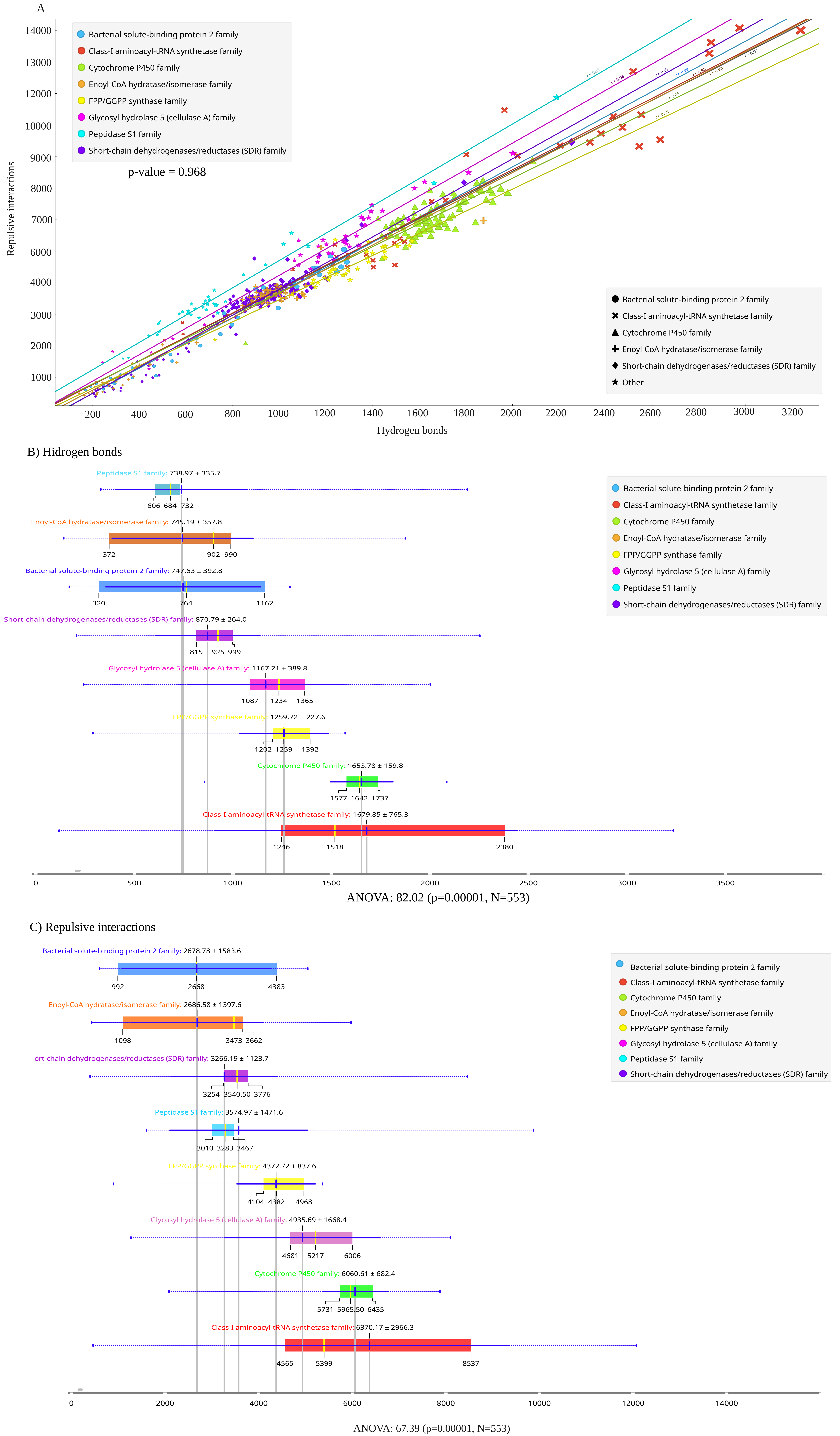
